# Targeting multidrug-resistant bacteria with genetic-information-free protein-only phages

**DOI:** 10.1101/2025.10.13.682247

**Authors:** Jeanette Batres, Branden Hunter, Hyunjin Shim

## Abstract

Bacteriophages offer advantages over small-molecule antibiotics, including host specificity and general compatibility with the human phagenome. However, their evolvability as replicating biological entities introduces therapeutic unpredictability and risks of phage-bacteria co-evolution. Here, we retain the targeting benefits of phages while avoiding genetic replication by engineering genetic-information-free, protein-only phages (POPs). These genome-free particles self-assemble in a cell-free protein synthesis system from modular, de novo gene fragments encoding only structural and antimicrobial proteins. Using *Enterobacteria phage* T7 and its susceptible bacterial host as a model, we test the hypothesis that POPs stochastically encapsulate small antimicrobial proteins during self-assembly and deliver them into bacteria during adsorption via an ejectome-mediated injection mechanism. A computational survey of T7 small proteins revealed early and mid-genome enrichments of hypothetical proteins and capsid volume sufficient to accommodate multiple small proteins in the absence of the ∼40-kb genome. In time-series antimicrobial susceptibility assays (48-72 h), POPs produced initial growth inhibition comparable to wild-type T7 at the highest doses with a linear dose-effect relationship and a minimum inhibitory concentration-like threshold. These results establish the feasibility of genetic-information-free POPs as protein-based antimicrobials that couple phage receptor specificity with minimal biosafety risks, supporting the development of more stable and predictable phage-inspired therapeutics.

## Introduction

Antimicrobial resistance (AMR) is a growing global health crisis, causing substantial mortality despite the widespread use of antibiotics and straining healthcare systems and markets (1). Beyond socioeconomic barriers to antibiotic development, previous reports reveal the rapid emergence of resistance to small-molecule drugs soon after deployment (2). Consequently, there is renewed interest in alternative antimicrobial strategies, including bacteriophages (phages) (3). Phages can selectively target pathogenic bacteria without affecting patient microbiomes, but their status as evolving biological entities introduces unpredictability, such as phage-bacteria co-evolution, and other unexpected outcomes that may complicate therapy (4,5).

Furthermore, bacteria have diverse anti-phage systems - at least as varied as canonical antibiotic-resistance mechanisms. Well-characterized systems include restriction-modification (RM) and CRISPR-Cas (6,7). More recently described defenses span cyclic-oligonucleotide signaling and abortive infection (e.g. CBASS, Thoeris, Abi), nucleotide depletion (e.g. dGTPases, AvcID), toxin-antitoxin and RNA/DNA nucleases (e.g. ToxIN, RnlAB, AVAST/Avs, DarTG), and cellular pathway disruption, among others (8).

Despite this anti-phage arsenal, phages routinely overcome host defenses with compact genomes. With genome sizes typically less than 50 kb - though huge phages exist in specific environments (9) -phages encode streamlined genetic programs with minimal noncoding DNA. Genomes are typically organized into early, middle, and late modules: early genes subvert host defenses and redirect cellular processes; middle genes drive transcription and replication; late genes specify virion assembly and lysis (Figure 1).

**Figure 1:**
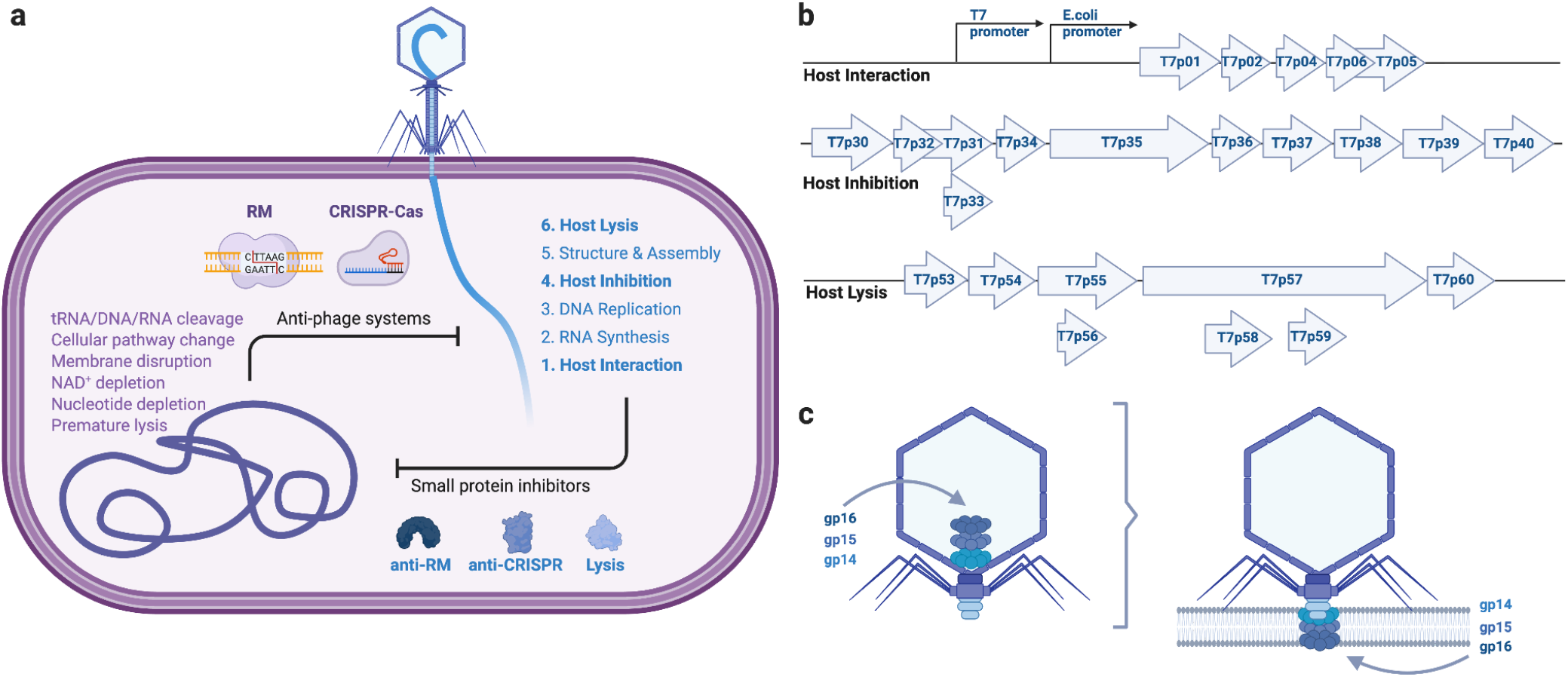
Schematic overview of phage infection and engineering strategy. (a) Host anti-phage defense systems and phage counter-defense factors across the infection cycle. (b) Design of genetic-information-free, protein-only phages (POPs) as modular ∼5 kb gene fragments encoding host interaction, host inhibition, and host lysis for cell-free protein synthesis. (c) Model of the T7 ejectosome. Before infection, gp14, gp15, and gp16 reside within the capsid. Upon adsorption, these subunits translocate in a (partially) unfolded state through the portal-tail complex, then reassemble into a transmembrane pore (ejectosome) that traverses the inner membrane of the host.

Within these modules, phage genomes frequently harbor clusters of small, often uncharacterized proteins. Many are plausible host-inhibition factors acting on pathways beyond nucleic-acid cleavage (e.g. metabolism, membranes, proteostasis). Small proteins are effective inhibitors because their compact, adaptable folds and frequent disorder enable diverse target engagement. Accordingly, they are often poorly captured by current structure predictors (10), which return low-confidence models for short or intrinsically disordered sequences. We hypothesize that a substantial fraction of these hypothetical small proteins exert bacteriostatic or bactericidal effects and together constitute an underexplored repertoire of protein antimicrobials.

Phage infection begins with adsorption to specific receptors and delivery of the viral genome by a tail apparatus that functions as a nanoscale injection machine. Tails can be short and noncontractile, long and contractile, or noncontractile, depending on the phage lineage (10). In *Enterobacteria phage* T7 (family *Podoviridae*), a short, noncontractile tail initiates release of internal virion proteins (gp14, gp15, gp16) that assemble an “ejectosome” channel across the cell envelope (11,12), enabling DNA translocation into the cytoplasm (Figure 1c). This step appears to be driven by conformational energy stored in the virion prior to access to cellular energy sources (13,14).

Building on the T7 ejectosome mechanism, we examined whether phage particles could be engineered to carry and eject non-genetic cargo. Prior work indicates that phage particles can act as nanoreactors and can be repurposed to transport small molecules (15,16). Here, we engineer protein-only phages (POPs) - genetic information-free systems designed to package and eject small antimicrobial proteins (AMPs) upon adsorption. Using de novo gene synthesis and cell-free protein synthesis, we assemble T7-derived capsids that stochastically co-encapsulate small antimicrobial proteins during self-assembly. Upon host adsorption, the virion’s built-in conformational energy (17) is repurposed to eject protein cargo - rather than genomic DNA - into the cytoplasm with receptor-defined specificity. Because POPs lack replicative capacity, they avoid phage-bacteria co-evolution during treatment and reduce the unpredictability associated with replicating biological agents. Derived from natural phage components yet devoid of genetic material, POPs may also mitigate certain regulatory and biosafety concerns related to genetic evolution or modification.

We evaluate these genetic-information-free, AMP-loaded POPs using transmission electron microscopy (TEM), dynamic light scattering (DLS), antimicrobial susceptibility testing (AST), plaque assays, and we analyze their small-protein cargo using structure prediction and biophysical characterization. Together, these results support a platform for delivering diverse antimicrobial proteins with the specificity of phage adsorption but without phage replication.

## Results

### Computational analysis of phage-derived small antimicrobial proteins

In the *Enterobacteria phage* T7 genome, genes are arranged in functional modules that track the infection cycle (Figure 1a). Small, poorly annotated genes are enriched at the 5′ end (T7p01-T7p06) and in a mid-genomic cluster (T7p30-T7p40) (Figure 1b). Because early phage gene products must first neutralize host defenses, we infer that the 5′ cluster likely encodes inhibitory effectors, consistent with the ocr-like anti-restriction factor (Table S1). We refer to this set (T7p01-T7p06) as antimicrobial by interaction (i.e. early host-targeting effectors). The mid-genomic cluster contains replication genes (e.g. polymerase) before another module of small inhibitors, such as a Gp5.9-like RecBCD inhibitor; we designate the mid-genome cluster of hypothetical proteins (T7p30-T7p40) as antimicrobial by inhibition (i.e. mid host-targeting effectors). Late genes specify virion assembly, with lytic functions (holin/spanins; T7p53-T7p56) embedded within the structural module.

Because experimental structures for most T7 hypothetical proteins are unavailable, we modeled the small proteins with AlphaFold (Figure 2a). Predicted template modeling (pTM) scores for many small proteins were <0.5, below confidence thresholds for high-quality predictions (Figure 2b). This mirrors prior observations that very short or intrinsically disordered phage proteins are under-represented in structure-prediction training sets (18). Low-confidence predictions were particularly frequent in the host-interaction and host-inhibition sets, which exhibited broad pTM distributions (mean ∼0.5 and ∼0.7, respectively). For each set, we show the model with the lowest pTM alongside the top-ranked model and its error profile (Figure 2c). These proteins typically had pLDDT < 70 and adopted rod-like or partially disordered conformations.

**Figure 2:**
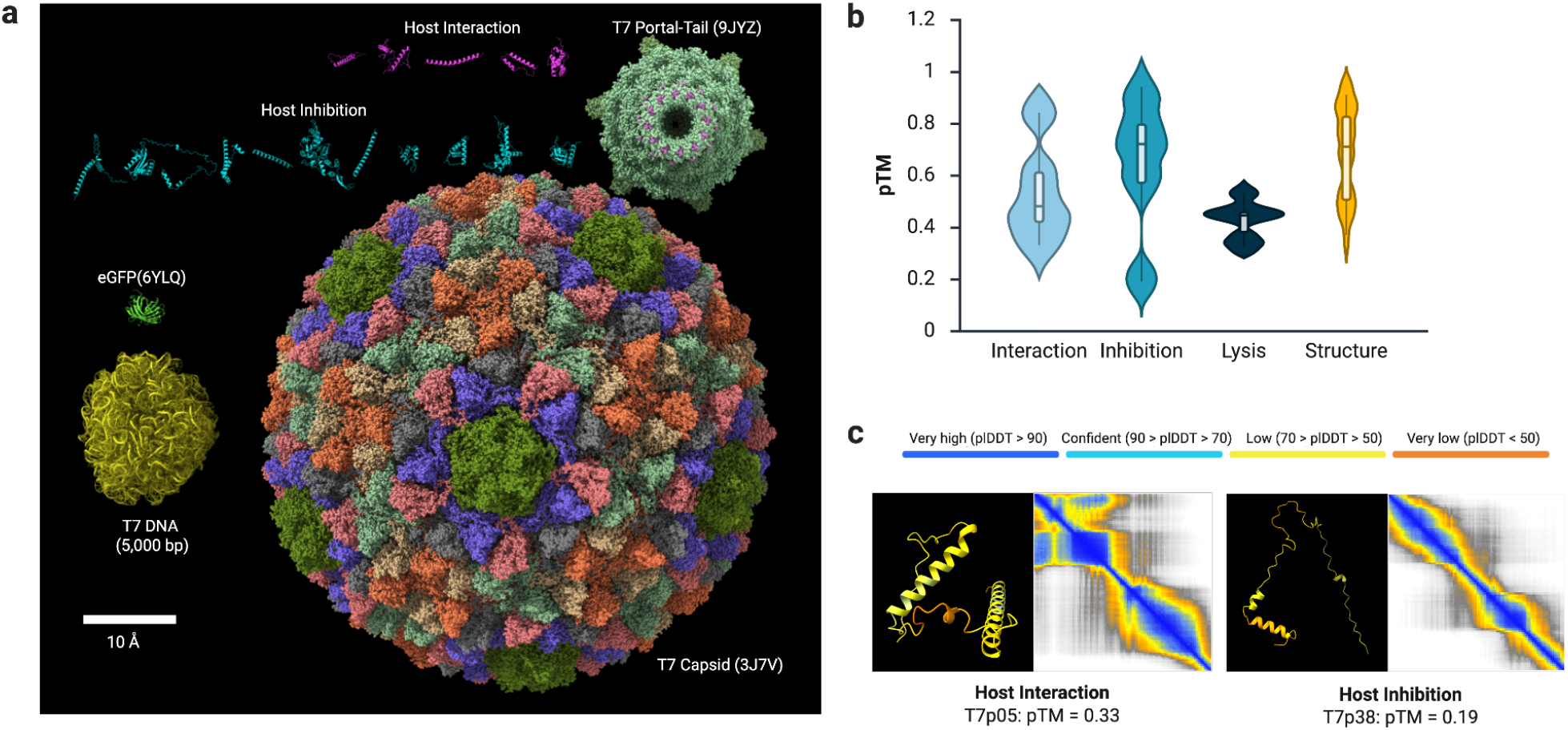
Phage-derived small antimicrobial proteins: scale and structural predictions. (a) Size and model comparison of the T7 capsid, a 5-kb DNA segment, representative phage-derived small proteins, and eGFP. (b) Distribution of AlphaFold pTM scores stratified by functional set in the T7 genome. (c) Representative models and confidence metrics for small antimicrobial proteins with the lowest pTM scores in each set (color-mapped by pLDDT).

We compared the sizes of small proteins from the interaction/inhibition sets with the T7 capsid (PDB: 3J7V), the portal-tail complex (PDB: 9JYZ), and a 5 kb DNA segment representative of the genome (total ∼40 kb) (Figure 2a). The capsid interior (∼60 nm diameter) readily accommodates numerous small proteins if no genomic DNA is enclosed, and portal dimensions are compatible with transit of small proteins, either natively folded or partially unfolded, analogous to ejectosome-mediated translocation during infection (Figure 1c). These geometric considerations support the hypothesis that capsids can serve as carriers for small-protein cargo, with portal-tail dynamics providing the mechanical drive for ejection (11,12).

### Experimental evaluation of phage-derived small antimicrobial proteins

As an exemplar for protein-only phages, we expressed the ocr-like anti-restriction protein (T7p01) fused to the N- and C-terminally truncated version of the reporter eGFP (deGFP) (19) to assess cell-free production and phage-mediated delivery. Visual comparison to a deGFP control plasmid indicated robust fluorescence after 16 h (≥ ∼400 μg/mL by the manufacturer’s fluorescence strip; Figure S1). NanoDrop A280 readings yielded higher apparent concentrations, as expected given contributions from cell-free protein synthesis (CFPS) components (RNA polymerase, ribosomes, auxiliary enzymes) (Table S2). Native PAGE showed a dominant band migrating between the 66 kDa and 20 kDa markers, consistent with the summed masses of deGFP (∼27 kDa) and T7p01 (∼14 kDa), alongside higher-mass bands attributable to CFPS background (Figure S1).

To test whether phage particles can carry small proteins, we co-expressed deGFP with the T7 structural genes in the CFPS system. The experimental structure of eGFP (PDB: 6YLQ) is comparable in size to other small proteins of interest (Figure 2a). Samples were desalted with 40-kDa-cutoff spin columns to deplete free deGFP and small molecules, which reduced background fluorescence. Upon exposure of *Escherichia coli* to the resulting preparations, fluorescence microscopy revealed faint but detectable cellular GFP signal within ∼20 min (phase-contrast and fluorescence modes; Figure S1), consistent with delivery of protein cargo associated with phage particles.

### Engineering protein-only phages with small antimicrobial proteins

The T7 genome was partitioned into ∼5,000-bp fragments spanning interaction, inhibition, structure, and lysis gene modules and rebooted using a cell-free transcription-translation system (Table S1). This modular architecture permits selective inclusion of gene sets while abolishing replicative capacity. De novo synthesis yielded high nucleic-acid amounts with acceptable purity (Table 1).

**Table 1:**
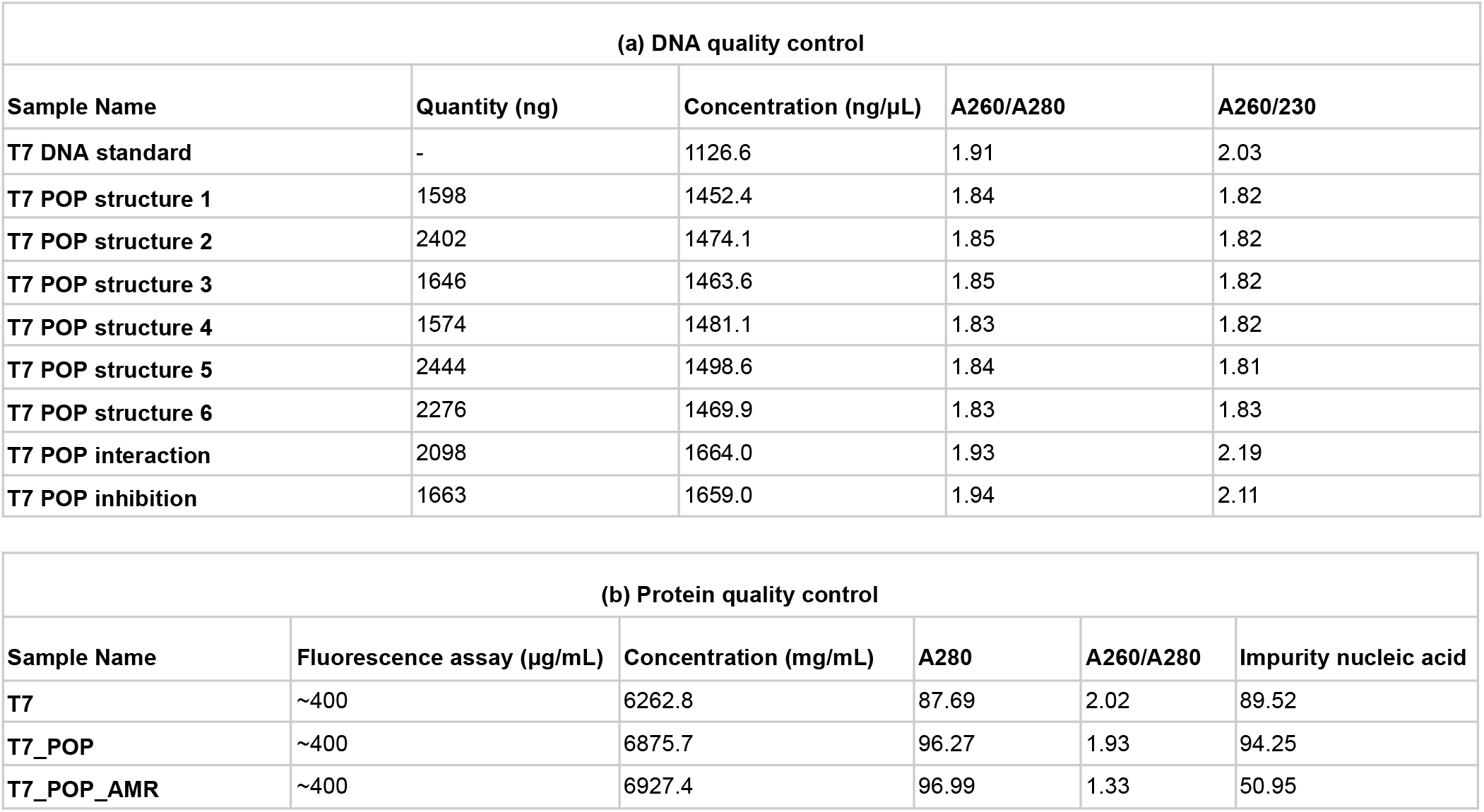
Quality control of DNA and protein during phage rebooting. (a) Spectrophotometric assessments of de novo-synthesized modular DNA fragments used for rebooting: absorbance at 260 nm (A260), 230 nm (A230), and 280 nm (A280), with corresponding concentrations and purity metrics. (b) Spectrophotometric assessments of samples after cell-free protein synthesis: absorbance at 280 nm (A280) and 260 nm (A260) for approximate protein content and nucleic-acid carryover, respectively, reported alongside estimated concentrations from fluorescence assays.

To confirm synthesis accuracy, we sequenced a subset of small-protein constructs (Table S3). For the ocr-like anti-restriction gene (T7p01), reads showed a mean Q score of 17.3 and a median read length of 555 bp (construct length: 554 bp). Alignment to the T7 reference produced a mean read-level identity >99.7%, as visualized in IGV; a minority of shorter reads mapped partially (Figure S2).

Following in vitro protein expression (Table S4), we assessed particle size, morphology, and concentration. First, we used Dynamic Light Scattering (DLS) to estimate the size distribution of T7 phage particles (Table S5). Three technical replicates indicated particles with a characteristic hydrodynamic radius (R_h) of ∼50 nm in both wild-type T7 preparations and engineered protein-only phages (POPs) (Figure 3). It is notable that size distributions for POPs were narrower than for wild-type, which displayed greater polydispersity (Figure S3).

**Figure 3:**
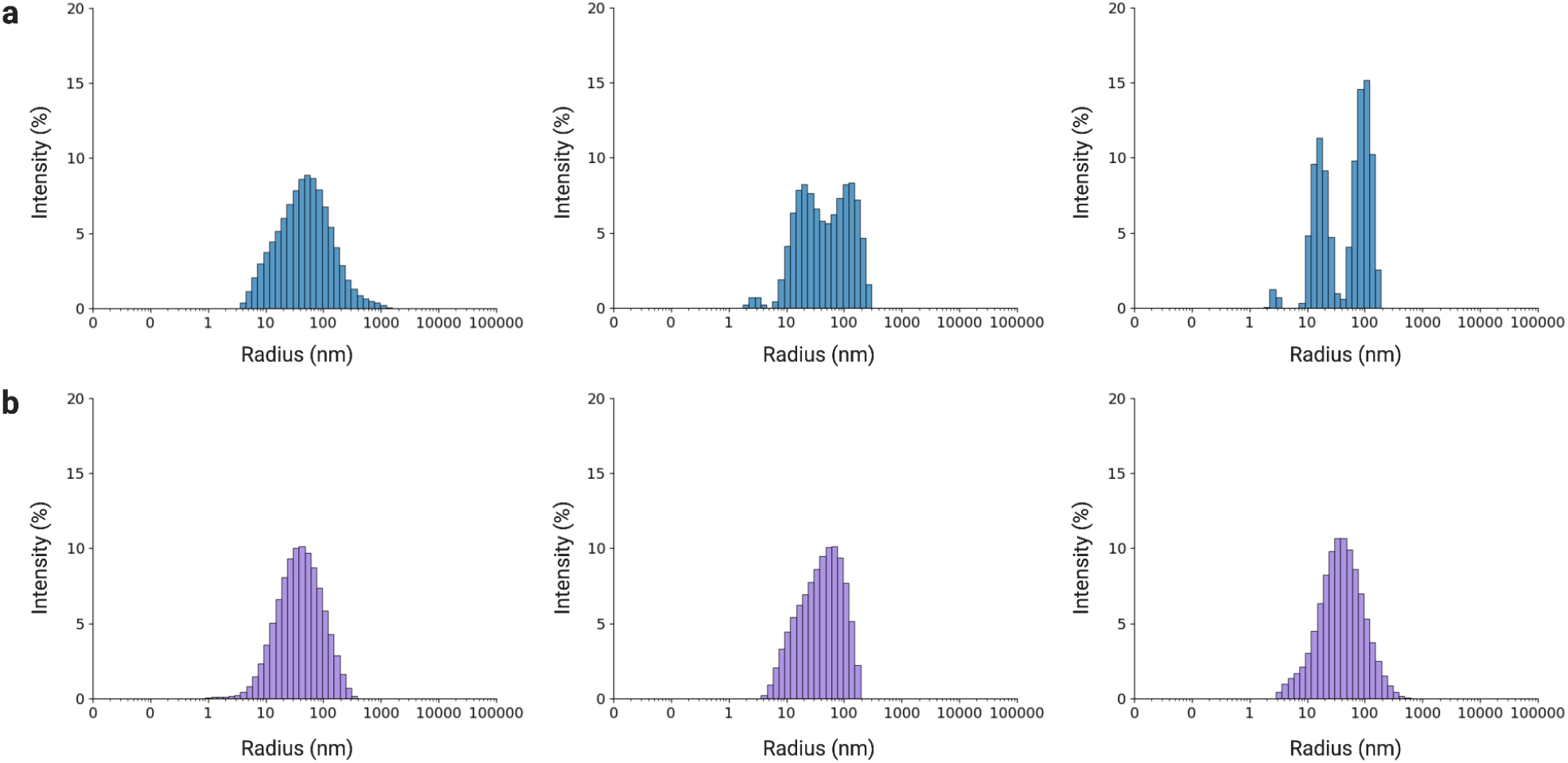
Dynamic light scattering of wild-type and engineered phage particles. (a) Wild-type T7: intensity-weighted hydrodynamic size distributions from three technical replicates. (b) POPs: intensity-weighted hydrodynamic size distributions from three technical replicates. Identical acquisition and analysis settings were used across conditions.

Second, we used Transmission Electron Microscopy (TEM) to visualize rebooted phage particles. After purification and negative staining, TEM micrographs (100 nm scale bars) revealed abundant icosahedral heads <100 nm across all phage preparations, consistent with the 60 nm T7 capsid diameter [36] (Figure S4). Short, noncontractile tails typical of podoviruses [35] of genome-free phage particles were not resolved, as reported in some TEM studies (20,21). Background residuals likely reflect remaining CFPS components. Collectively, these images support the structural integrity of POPs assembled by de novo gene synthesis and cell-free protein synthesis.

Third, we measured the concentration of proteins that contain amino acids such as aromatic residues and disulfides that exhibit absorbance at 280 nm (22). A280 measurements indicated high apparent protein concentrations in all rebooted samples (> 6,000 mg/mL) with detectable nucleic acid background (Table 1). As shown with a deGFP control, A280 overestimates due to CFPS constituents; we therefore interpret POP concentrations as ∼400 μg/mL, similar to the previous estimate (Figure S1).

Last, we used a plaque assay of wild-type T7 phages to estimate the concentration of POP during the antimicrobial susceptibility testing. As expected, POPs yielded no plaques in PFU assays, whereas wild-type T7 titers yielded dense plaque-forming units at the two highest doses (see below for details). These benchmarks guided dose selection for antimicrobial susceptibility testing.

### Antimicrobial susceptibility testing of engineered protein-only phages

We quantified growth inhibition of *Escherichia coli* (Migula) Castellani and Chalmers, reported as multidrug-resistant and susceptible to T7 (23), by time-series OD600 monitoring in 96-well plates over 48h (read every 20 min; three technical replicates; Table S6). Wild-type T7 served as a positive control; HPLC-grade water served as a negative control (Figure 4a).

**Figure 4:**
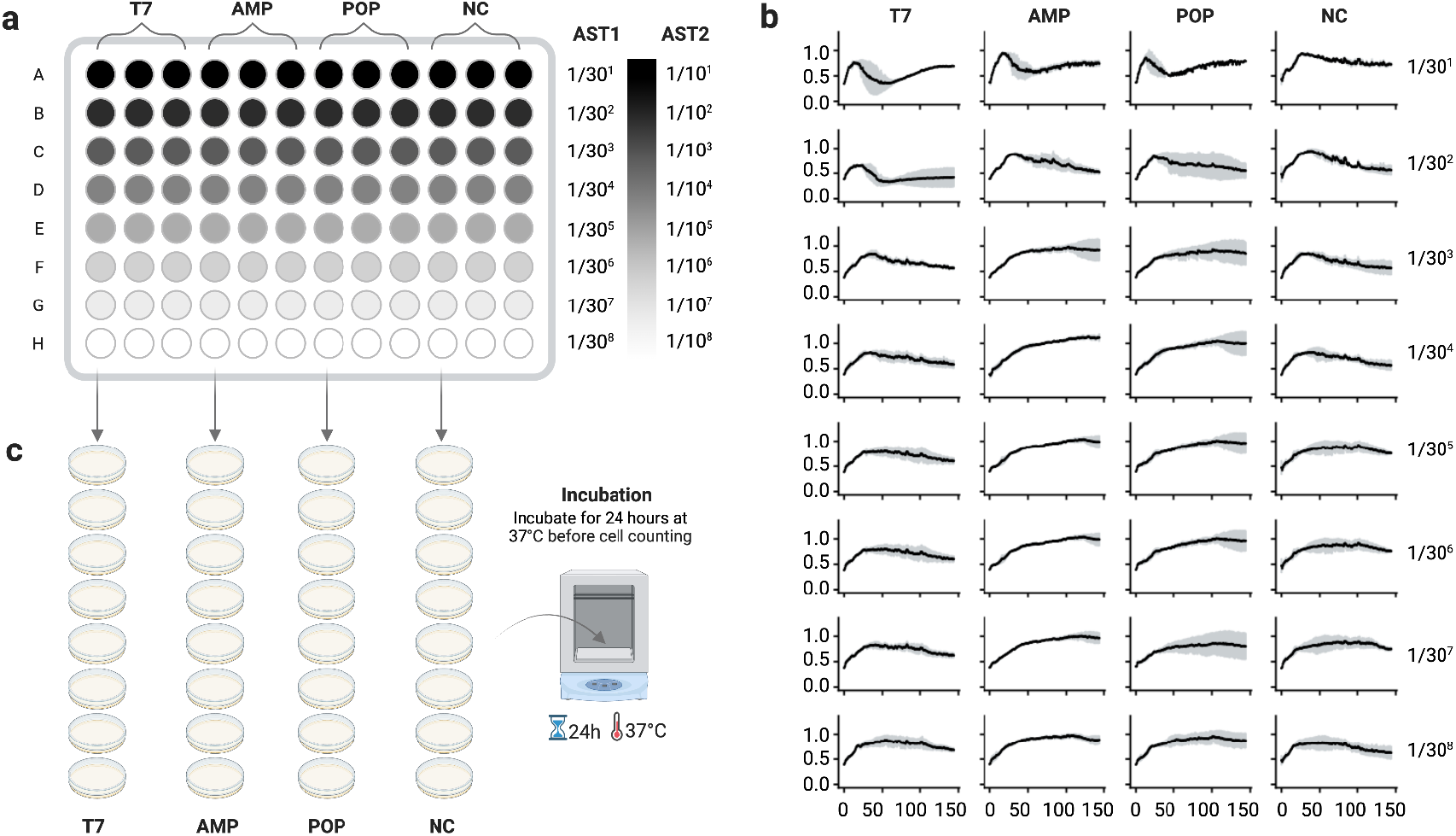
Antimicrobial susceptibility testing of protein-only phages against *E. coli*. (a) Plate map and dilution scheme. Each well contains 120 µL of *E. coli*; 4 µL of test sample is added to the first row, and 30-fold serial dilutions are performed down each column. This scheme is repeated for three technical replicates. (b) Time-series OD600 measurements acquired every 20 min for 48 h at 37°C. Points denote the mean of three technical replicates; grey error bars indicate SD. Rows correspond to successive 30-fold dilutions. Conditions: wild-type T7 (positive control, PC), AMP-loaded POPs (AMP), POPs, and HPLC-grade water (negative control, NC). (c) Plaque assay on an *Escherichia coli* lawn. For each plate, 120 µL of cells were mixed with 4 µL of test sample, incubated for 10 min at room temperature, spread on preheated LB agar plates, and incubated inverted at 37 °C overnight.

In the highest-dose row, all phage conditions of wild-type T7, AMP-loaded POPs, and POPs showed strong initial suppression relative to the negative control (Figure 4b). At 12 h, the OD600 values for the T7, AMP, and POP samples were 49%, 76%, and 65%, respectively, relative to the OD600 value for the negative control (Table 2). Inhibition waned after ∼24 h across conditions, including wild-type T7, consistent with the emergence of phage-tolerant/resistant subpopulations. For engineered POPs, this time-dependent reduction in effectiveness may be due to the dilution of effective particle-to-cell ratios. The negative-control trajectory followed the expected exponential-stationary-death phases (24).

**Table 2:** Antimicrobial susceptibility testing of negative controls, positive controls, and protein-only phages Summary of three replicates of OD600 readings from a microplate reader every 20 minutes. (a) AST assay for 48 hours with a dilution factor of 30. (b) AST assay for 72 hours with a higher initial dose and a dilution factor of 10.

| (a) AST1 |  |  |  |  |  |
| --- | --- | --- | --- | --- | --- |
|  | OD600 | T7<br>(mean±stdev) | AMP<br>(mean±stdev) | POP<br>(mean±stdev) | NC<br>(mean±stdev) |
| Dilution<br>factor<br>=30<sup>1</sup> | t=0h | 0.346±0.011 | 0.375±0.029 | 0.371±0.005 | 0.436±0.116 |
|  | t=12h | 0.448±0.313 | 0.696±0.227 | 0.599±0.092 | 0.919±0.033 |
|  | t=24h | 0.418±0.033 | 0.638±0.046 | 0.615±0.026 | 0.839±0.011 |
|  | t=36h | 0.635±0.004 | 0.744±0.057 | 0.779±0.019 | 0.738±0.044 |
|  | t=48h | 0.691±0.011 | 0.743±0.061 | 0.789±0.010 | 0.726±0.038 |
| Dilution<br>factor<br>=30<sup>2</sup> | t=0h | 0.385±0.008 | 0.391±0.004 | 0.386±0.012 | 0.485±0.164 |
|  | t=12h | 0.509±0.170 | 0.880±0.048 | 0.805±0.116 | 0.938±0.050 |
|  | t=24h | 0.362±0.074 | 0.769±0.133 | 0.690±0.229 | 0.814±0.057 |
|  | t=36h | 0.404±0.169 | 0.628±0.112 | 0.635±0.268 | 0.606±0.058 |
|  | t=48h | 0.421±0.195 | 0.521±0.049 | 0.551±0.152 | 0.562±0.104 |
| Dilution<br>factor<br>=30<sup>3</sup> | t=0h | 0.380±0.001 | 0.383±0.002 | 0.384±0.012 | 0.374±0.015 |
|  | t=12h | 0.843±0.077 | 0.831±0.014 | 0.809±0.044 | 0.846±0.038 |
|  | t=24h | 0.719±0.041 | 0.934±0.038 | 0.882±0.102 | 0.734±0.076 |
|  | t=36h | 0.636±0.036 | 0.968±0.067 | 0.905±0.184 | 0.614±0.097 |
|  | t=48h | 0.569±0.037 | 0.921±0.217 | 0.850±0.233 | 0.570±0.143 |

|  | (b) AST2 |  |  |  |  |
| --- | --- | --- | --- | --- | --- |
|  | OD600 | T7<br>(mean±stdev) | AMP<br>(mean±stdev) | POP<br>(mean±stdev) | NC<br>(mean±stdev) |
| Dilution<br>factor<br>=10<sup>1</sup> | t=0h | 0.289±0.042 | 0.272±0.013 | 0.293±0.024 | 0.485±0.371 |
|  | t=12h | 0.486±0.083 | 0.435±0.041 | 0.441±0.021 | 0.687±0.038 |
|  | t=24h | 0.613±0.077 | 0.610±0.085 | 0.610±0.054 | 0.738±0.137 |
|  | t=36h | 0.773±0.018 | 0.759±0.062 | 0.767±0.029 | 0.746±0.116 |
|  | t=48h | 0.842±0.033 | 0.917±0.027 | 0.856±0.035 | 0.720±0.071 |
|  | t=60h | 0.868±0.048 | 0.904±0.016 | 0.891±0.059 | 0.690±0.013 |
|  | t=72h | 0.885±0.046 | 0.934±0.030 | 0.892±0.040 | 0.702±0.004 |
| Dilution<br>factor<br>=10<sup>2</sup> | t=0h | 0.303±0.019 | 0.302±0.020 | 0.303±0.011 | 0.355±0.105 |
|  | t=12h | 0.543±0.131 | 0.511±0.050 | 0.448±0.049 | 0.659±0.066 |
|  | t=24h | 0.544±0.097 | 0.516±0.023 | 0.488±0.005 | 0.597±0.019 |
|  | t=36h | 0.562±0.105 | 0.483±0.022 | 0.486±0.004 | 0.581±0.032 |
|  | t=48h | 0.561±0.088 | 0.519±0.027 | 0.496±0.017 | 0.542±0.059 |
|  | t=60h | 0.554±0.075 | 0.527±0.021 | 0.506±0.020 | 0.533±0.078 |
|  | t=72h | 0.591±0.103 | 0.560±0.017 | 0.530±0.029 | 0.577±0.113 |
| Dilution<br>factor<br>=10<sup>3</sup> | t=0h | 0.307±0.020 | 0.305±0.028 | 0.550±0.443 | 0.475±0.298 |
|  | t=12h | 0.598±0.065 | 0.700±0.034 | 0.683±0.056 | 0.635±0.038 |
|  | t=24h | 0.618±0.048 | 0.843±0.033 | 0.776±0.104 | 0.600±0.026 |
|  | t=36h | 0.584±0.052 | 0.895±0.069 | 0.660±0.173 | 0.585±0.043 |
|  | t=48h | 0.537±0.030 | 0.880±0.044 | 0.578±0.186 | 0.546±0.083 |
|  | t=60h | 0.528±0.024 | 0.833±0.104 | 0.536±0.207 | 0.540±0.106 |
|  | t=72h | 0.522±0.005 | 0.861±0.079 | 0.564±0.191 | 0.550±0.108 |

As the dose decreased across rows, inhibition diminished. For wild-type T7, the minimum inhibitory concentration (MIC)-like threshold occurred between rows 2-3 (dilution factor = 30^3^), after which trajectories resembled the negative control. For POPs, the threshold was between rows 1-2 (dilution factor = 30^2^), indicating a higher effective dose requirement without replication. Across concentrations, AMP-loaded POPs exhibited smaller SD envelopes in grey error bars than POPs, suggesting reduced sample-to-sample variability.

PFU assays performed in parallel confirmed that engineered POPs without replicative capacity produced no plaques at any dilution, whereas wild-type T7 yielded countable plaques at higher doses (Figure 4c). The highest dose of wild-type T7 phages showed 1,224±12 PFUs, and the second highest dose showed 212±2 PFUs (Figure S5). From the third highest dose, no PFU was observed, which is consistent with the results from the antimicrobial susceptibility testing. The concentration of wild-type T7 phages was calculated to be 3.06×10^5^ PFU/mL and 1.59×10^6^ PFU/mL for the highest and second-highest doses, respectively. The high plaque density observed at the highest dose likely led to undercounting of plaques, resulting in an underestimation of the true titer beyond the confidence range. Although POPs lack replicative capacity and therefore do not yield PFUs, orthogonal particle-based readouts (DLS, TEM, A280) from our rebooted preparations suggest that engineered particles assembled by de novo gene synthesis and cell-free protein synthesis are present at concentrations comparable to wild-type preparations generated under matched conditions.

### Higher-dose, extended assay toward bactericidal endpoints

Because growth recovered after ∼24 h in the MIC assay, we repeated the experiment with increased starting doses and lower dilution factors, and extended monitoring to 72 h (three technical replicates; Tables S7-8). The highest-dose row again showed strong early inhibition for all phage conditions, followed by loss of effect by ∼24 h (Figure 5). At 12 h, the OD600 values for the T7, AMP, and POP samples at the highest dose were 70%, 63%, and 64%, respectively, relative to the OD600 value for the negative control (Table 2). Dose-response trends persisted: inhibition faded between rows 2-3 (dilution factor = 10^3^) for both wild-type and POPs. Together with the previous MIC assay, these data are consistent with an approximately linear dose-effect relationship for POPs over the tested range, whose effectiveness increases with increasing dosage (Figure 5b). However, wild-type T7 showed less predictable long-term dynamics as a replicating agent, with less reproducible trajectories (larger SD envelopes shown as grey error bars) and an unclear dose-effect relationship. Under the present conditions, bactericidal endpoints were not sustained for either wild-type T7 or POPs used as sole agents, indicating that higher effective particle numbers and/or combination strategies may be required to achieve minimum bactericidal concentrations.

**Figure 5:**
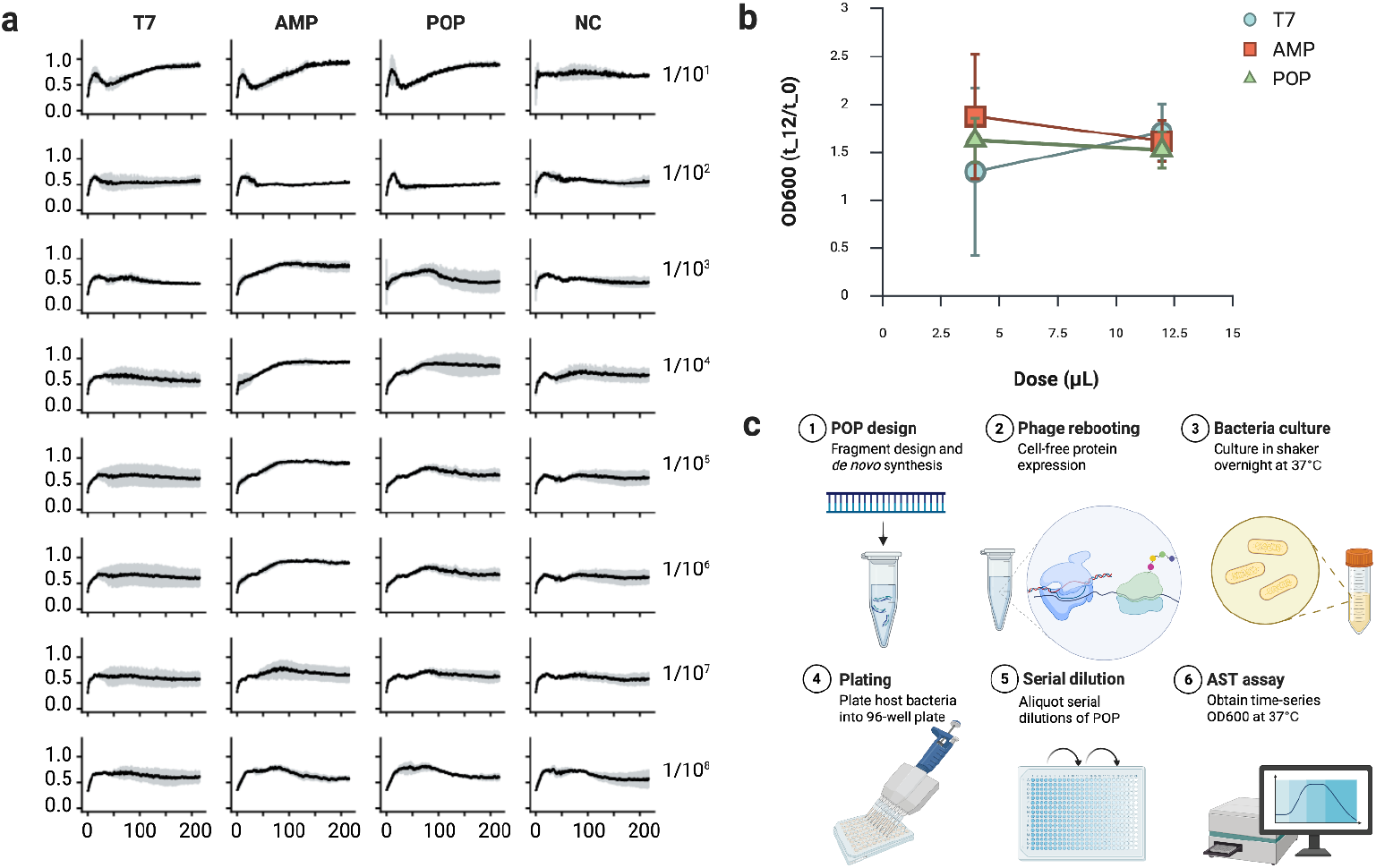
Extended antimicrobial susceptibility testing of protein-only phages (higher dose, extended monitoring) (a) Time-series OD600 measurements acquired every 20 min for 72 h at 37°C. Points denote the mean of three technical replicates; grey error bars indicate SD. Rows correspond to successive 10-fold dilutions. Conditions: wild-type T7 (positive control, PC), AMP-loaded POPs (AMP), POPs, and HPLC-grade water (negative control, NC). (b) Dose-effect relationship of wild-type T7 phages and engineered POPs based on antimicrobial susceptibility testing (AST) assays, shown as the relative increase in OD600 at 12 hours compared to the initial OD600. (c) Workflow for POP generation and testing: 1. Design and de novo synthesize modular linear gene fragments encoding antimicrobial and structural proteins. 2: Reboot fragments in a cell-free protein synthesis system to generate POPs. 3: Culture the host bacteria in LB broth at 37°C, 220 rpm overnight. 4: Dispense cells into 96-well plates. 5: Apply serial dilutions of POPs across the plate. 6: assess inhibitory activity by time-series monitoring of OD600.

## Discussion

Proteins are the primary functional hardware of biology, capable of extraordinary chemical and structural diversity arising from just 20 amino acids and their post-translational modifications (25). Many phage counter-defense strategies are protein-encoded - often by short, compact effectors that act on host nucleases, membranes, metabolism, or transcriptional machinery (26,27). Consistent with prior observations, our computational analysis highlights the prevalence of small, poorly characterized proteins in early and mid T7 phage genomic modules, where counter-defense and host takeover are initiated to redirect host resources for phage replication. Structure prediction for these sequences reveals the unique forms and the known limitations of current predictors for short and intrinsically disordered proteins (18).

We introduce protein-only phages (POPs) - genetic-information-free phage particles intended to package and eject small antimicrobial proteins upon adsorption. Using de novo gene synthesis and cell-free protein synthesis, we generated modular, non-replicative assemblies and verified particle formation by DLS and TEM. In antimicrobial susceptibility testing, POPs produced a strong early suppression of *E. coli* comparable to wild-type T7 at the high dose. Furthermore, they displayed a more predictable, approximately linear dose-effect relationship across the tested range, unlike replicating wild-type T7 phages. Under the present conditions, neither wild-type T7 nor POPs achieved bactericidal activity, with growth recovery after ∼24 hours.

The T7 injection apparatus (gp14/gp15/gp16) deploys conformation energy stored in the virion to assemble an ejectosome that spans the bacterial cell envelope, enabling genome translocation (11,12). By analogy, our results with a fluorescent protein provide indirect evidence that protein cargo associated with phage particles can enter the cytoplasm. The observed linear dose-effect for POPs is consistent with a stoichiometric delivery model, in contrast to the complex, time-dependent dynamics of replicating wild-type phages. Two mechanistic variables merit attention: stochastic cargo loading (copy number, localization, and stability within capsids) and translocation efficiency through the portal-tail complex, which may depend on cargo size, folding state, charge, and interactions with internal virion proteins.

Because POPs lack intact nucleic acids and do not replicate, risks associated with lysogeny, transduction, and phage-bacteria co-evolution are intrinsically eliminated. This makes POPs more attractive alternative antimicrobials, given a similar effectiveness in bacterial growth inhibition to wild-type phages. Furthermore, POPs are expected to be safe for the human body, as they are derived from natural phages and a human phageome has recently been described (28–30). However, capsid proteins and proteinaceous cargos are still biological macromolecules; immunogenicity, biodistribution, and clearance will require rigorous preclinical evaluation (31). Likewise, diverse phages with receptor specificity that benefits targeting should be investigated for clinically relevant pathogens, such as those on the WHO bacterial priority pathogens list (32).

Several aspects of our study need further validation before preclinical evaluation. First, delivery of small proteins by phage particles was supported by bacterial growth phenotypes and microscopy, but not yet demonstrated by direct intracellular quantification. Second, a direct measurement of POP dose and cargo copy number may be necessary for preclinical studies, potentially through particle-centric assays and targeted proteomics. Third, MIC-format assays showed regrowth of bacteria across conditions, suggesting adsorption-resistant subpopulations, proteolytic degradation of cargo, or insufficient particle-to-cell ratios limit durability.

Next steps for this genetic-information-free system include combining POPs with antibiotics or phages to achieve durable killing while preserving dose-response predictability. Additional protein cargos, such as AI-designed antimicrobial peptides, can be explored using advances in generative AI, de novo gene synthesis, and cell-free protein synthesis. As all experiments were performed in a single *E. coli* model, evaluation across broader panels of key pathogens is needed to generalize performance. Stability, formulation, and pharmacokinetic studies will also be essential for translational development.

POPs leverage the specificity of phage adsorption and the mechanical work of the ejectosome to deliver antimicrobial proteins without genetic material or modification. While our data establish feasibility and reveal dose-response behavior, achieving sustained bactericidal activity will likely require improved cargo loading, enhanced delivery efficiency, and combination therapy. With these advances, POPs could emerge as a programmable platform for protein-based antimicrobials targeted to drug-resistant bacteria.

## Methods

### Computational structure prediction of small antimicrobial proteins

Small-protein sequences of the NCBI Reference Sequence of *Enterobacteria phage* T7 (NC_001604.1) were compiled (Table S1) and categorized by locus cluster into “host interaction” (T7p01-T7p06), “host inhibition” (T7p30-T7p40), and “host lysis” (T7p53-T7p60). Structures were predicted using the AlphaFold Protein Structure Prediction server (EMBL-EBI; accessed on 2025/05/25), and the top-ranked model from each run was retained (33,34). Figures were generated in ChimeraX with per-residue plDDT mapped to the models, and summary statistics including plDDT (per-residue confidence), PAE (predicted aligned error), and pTM scores are reported for all predictions (Figure 2).

### Cell-free expression of small antimicrobial proteins

Two fusion constructs of ocr-like anti-restriction gene (T7p01) fused to deGFP at either the 5′ or 3′ terminus (5′-deGFP-Ocr and Ocr-deGFP-3′) were expressed alongside a deGFP control in a cell-free transcription-translation system (myTXTL Pro Kit, Daicel Arbor Biosciences, USA). As fluorescence tests, crude protein levels were first estimated visually with the manufacturer’s color strip (Figure S1). Protein concentrations were then measured by A280 on a NanoDrop Microvolume UV-Vis Spectrophotometer (Thermo Scientific, USA). The molecular weight for A280 processing was set to the instrument maximum (9,999 kPa), and the extinction coefficient (ε280) to 1,400×10^3^ M^−1^ cm^−1^ (Table S2).

Samples were analyzed by native PAGE (4-16% Bis-Tris; ThermoFisher, USA). Preparations used a native (non-reducing, non-denaturing) loading buffer and were maintained on ice until loading. NativeMark™ Unstained Protein Standard (ThermoFisher, USA) and the Ocr-deGFP-3′ were run in parallel to estimate apparent native masses. Following electrophoresis, gels were stained with Coomassie Blue R-250 (ThermoFisher, USA) and imaged on a GelDoc Imaging System (Bio-Rad, USA) to assess total protein (Figure S1). Band positions and relative intensities were quantified with identical exposure and analysis settings across conditions (ImageLab v4.1).

Fluorescence microscopy was used to monitor changes in cellular fluorescence in *Escherichia coli* ATCC® 8739™ (Microbiologics, USA) exposed to phages rebooted with deGFP proteins. Rebooted samples were filtered using Zeba Spin Desalting columns (cutoff: < 40-kDa proteins and < 2-kDa small molecules). Host cells and matched controls (cells alone, cells with wild-type T7 lacking deGFP, and cells with purified deGFP) were prepared in parallel. Aliquots (5 µL) were placed on poly-L-lysine-coated adhesive microscope slides (LabScientific, USA) and imaged on a fluorescence system equipped with a GFP-appropriate filter set under temperature-controlled conditions. Image acquisition parameters (focus, exposure, illumination) were held constant within experiments to enable quantitative comparison (Figure S1).

### Design and synthesis of protein-only phage constructs

The NCBI Reference Sequence of *Enterobacteria phage* T7 (NC_001604.1) served as the design template. Using RefSeq annotations, structural and antimicrobial open reading frames (ORFs) were selected and assembled into a linear gene fragment of ∼5000 base pairs, flanked by a T7 promoter and terminator and capped with provider-recommended universal adapters (Table S1). Custom de novo gene synthesis was performed according to the provider’s proprietary protocols (Twist Bioscience, USA) via high-throughput, silicon-based synthesis per provider protocols with a reported average error rate of 0.013%. On-platform assembly comprised annealing and PCR amplification of initial oligonucleotides on the semiconductor substrate, followed by enzymatic error correction. Upon receipt, each fragment was resuspended in 10 µL TE buffer (10 mM Tris-Cl, 0.5 mM EDTA, pH 9.0) and stored at −20 °C. Sequence identity and integrity were assessed by nanopore sequencing (Oxford Nanopore Technologies, UK) using Native Barcoding Kit (SQK-NBD114.24) and a MinION flow cell (R10.4.1). Reads were aligned to the T7 reference genome with minimap2 (35) and visualized in IGV (v2.19.5) to inspect coverage and mismatches (Figure S2).

### Cell-free transcription-translation of engineered phages

Phage rebooting and protein expression were performed using a cell-free transcription-translation system (myTXTL Pro Kit, Daicel Arbor Biosciences, USA), which provides both *Escherichia coli* and T7 RNA polymerases and enables expression without cloning, lysis, or purification (36,37). As a positive control, a commercially sourced bacteriophage T7 genomic DNA standard (Boca Scientific, USA) was used. Reaction components (Pro Master Mix, Pro Helper Plasmid, nuclease-free water, and T7 genomic DNA) were assembled according to the manufacturer’s instructions, with DNA supplied at the recommended 0.25 nM for T7 rebooting (Table S4). After incubation per kit guidelines, reactions were transferred to ice pending downstream analysis.

For protein-only phage constructs, structural and antimicrobial genes were expressed from synthesized linear DNA fragments (as described above). Reactions contained Pro Master Mix and Pro Helper Plasmid, with linear templates at the manufacturer-recommended 20 nM. Template inputs were calculated from the molar concentrations, and overall reaction compositions are summarized in Table S4. Following incubation per kit guidelines, reactions were transferred to ice pending downstream analyses.

Reaction products were quantified on a NanoDrop Microvolume UV-Vis Spectrophotometer. A280 measurements were collected using 1 µL of each sample, with the molecular-weight parameter set to 9,999 kPa and the extinction coefficient (ε280) set to 1,400×10^3^ M^−1^ cm^−1^. A260 was also recorded to estimate contributions from nucleic acids and other transcription-translation components (Table 1).

### Dynamic light scattering and transmission electron microscopy of rebooted T7 phages

Hydrodynamic size distributions were measured by Dynamic Light Scattering (DLS) on a DynaPro NanoStar II (Wyatt Technology, USA) equipped with a 658 nm laser and 90° optics. Rebooted phage preparations (4 µL) and matched buffer blanks were loaded into disposable cyclic-olefin-copolymer (COC) microcuvettes under temperature control per the manufacturer’s recommendations. Three technical replicates were acquired for each sample (Table S5).

Prior to imaging, phage preparations were purified by fast protein liquid chromatography (FPLC) on a Bio-Rad NGC Chromatography System to remove small proteins, nucleic acids, and buffer components that interfere with electron microscopy. A size-exclusion workflow was used to separate intact virions from lower-molecular-weight species under isocratic conditions. Sterile filtered phosphate-buffered saline (PBS; ThermoFisher, cat. J61196.AP) served as the equilibration, sample, and elution buffer.

For TEM sample preparation, carbon-coated 400-mesh grids (Ted Pella, USA) were used for negative staining. Freshly prepared phage sample (2 µL) was applied to the carbon side for 30 s, then washed with drops of 30 µL HPLC-grade water (Fisher Scientific, USA). Grids were immediately stained with 5 µL of 1% uranyl acetate for 30 s, blotted with Whatman paper, and air-dried overnight. Imaging was performed on a Talos F200C G2 at the Imaging and Microscopy Facility, University of California, Merced.

### Antimicrobial susceptibility testing of protein-only phages

A standard suspension of *Escherichia coli* ATCC® 8739™ (Microbiologics, USA) served as the host organism. An aliquot (10 µL) was inoculated into liquid LB broth overnight at 37 °C with agitation at 220 rpm. Initial and final cell densities were recorded by OD600 readings using a NanoDrop Microvolume UV-Vis Spectrophotometer. Cultured cells (120 µL per well) were dispensed into 96-well plates. Test samples (4 µL) were serially diluted from the first to the last row using a constant dilution factor of 30, with three technical replicates per condition (Figure 4). Plates were immediately loaded onto a SpectraMax iD3 (Molecular Devices, USA) and run at 37 °C for 48 hours with OD600 readings every 20 min after dosage (Table S6).

Engineered protein-only phages (POP and AMP-loaded POP) were tested by adding 4 µL of each sample to the first row containing 120 µL of cultured *E. coli*, then serially diluting across rows by transferring 4 µL of the mixture. For the positive control (PC), wild-type T7 phages rebooted from the standard phage DNA were added (4 µL) to the first row containing 120 µL of cultured *E. coli*, then serially diluted across rows by transferring 4 µL of the mixture. For the negative control (NC), HPLC-grade water (Fisher Scientific, USA) was added (4 µL) to the first row containing 120 µL of cultured *E. coli*, followed by the same serial dilution scheme.

Immediately after preparation, 120 µL of each test mixture was incubated for 10 min at room temperature for the plaque-forming unit (PFU) assay (38). Each test mixture was plated on LB agar plates (Thermo Fisher, USA) preheated to 37 °C, allowed to stand for 15 min at room temperature, and then incubated inverted at 37 °C overnight (Figure 4). Plates were imaged over a grid (Figure S5), and images were imported into OpenCFU (v3.9) and GoodNotes 5 (v6.6.16) for plaque counting (39). The automated plaque counting was checked manually by grid-based counts to obtain totals. PFU per milliliter was calculated as:

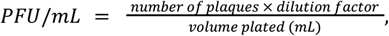

with the dilution factor taken from the serial dilution series (Figure 4).

### Extended antimicrobial susceptibility testing of protein-only phages

For a second antimicrobial susceptibility test (AST), we used a higher starting concentration of each test sample than in the first AST assay. A standard suspension of *Escherichia coli* ATCC® 8739™ (Microbiologics, USA) served as the test organism. An aliquot (12 µL) was inoculated into liquid LB broth and incubated overnight at 37 °C with agitation at 220 rpm. Initial and final cell densities were recorded by OD600 readings on a NanoDrop Microvolume UV-Vis Spectrophotometer. Cultured cells (120 µL per well) were dispensed into 96-well plates. Test samples were serially diluted from the first to the last row with a constant dilution factor of 10, with three technical replicates per condition (Figure 4). Plates were immediately loaded onto a SpectraMax iD3 (Molecular Devices, USA) and run at 37 °C for 72 hours with OD600 readings every 20 minutes after dosage (Table S8).

Engineered protein-only phages (POP and AMP-loaded POP) were tested by adding 12 µL of each sample to the first row containing 120 µL of cultured *E. coli*, then serially diluting across rows by transferring 12 µL of the mixture. For the positive control (PC), wild-type T7 phages rebooted from the standard phage DNA were added (12 µL) to the first row containing 120 µL of cultured *E. coli*, then serially diluted across rows by transferring 4 µL of the mixture. For the negative control (NC), HPLC-grade water (Fisher Scientific, USA) was added (12 µL) to the first row containing 120 µL of cultured *E. coli*, followed by the same serial dilution scheme.

## Declarations

### Ethics approval and consent to participate

Not Applicable

### Consent for publication

Not Applicable

### Availability of data and materials

Supplementary Tables (single workbook with multiple tabs). For data analysis, R v.3.6.0 and ggplot2 v.3.3.0 were used.

### Authors’ contributions

JB: data collection, analysis, and interpretation, manuscript preparation. BH: data collection, analysis, and interpretation, manuscript preparation. HS: study conception and design, data collection, analysis, and interpretation, manuscript preparation.

## Acknowledgments

We thank Masaki Uchida, Cory Brooks, Alejandro Calderon-Urrea at California State University, Fresno, and Kennedy Nguyen at the Imaging and Microscopy Facility at the University of California, Merced.

